# Actin monomers influence the interaction between *Xenopus* cyclase-associated protein 1 and actin filaments

**DOI:** 10.1101/2025.08.14.670363

**Authors:** Phuong Doan N. Nguyen, Hiroshi Abe, Shoichiro Ono, Noriyuki Kodera

## Abstract

Cyclase-associated protein (CAP) binds to both actin monomers and filaments and regulates multiple aspects of the actin dynamics including polymerization and depolymerization. CAP has been isolated from multiple species as a stable equimolar complex with actin monomer. However, functional significance of the CAP-actin complex is unknown. We previously demonstrated that native *Xenopus* cyclase-associated protein 1 (XCAP1) forms a 4:4 complex with actin. Here, we characterized how actin-free XCAP1 and the XCAP1-actin complex interact with actin filaments using high-speed atomic force microscopy and found that XCAP1-bound actin monomers influence dwell time and positional preference of XCAP1 on actin filaments. Actin-free XCAP1 bound to actin filaments transiently with a dwell time of ∼0.2 seconds. The XCAP1-actin complex also bound to actin filaments transiently but with a 3-to 5-fold longer dwell time than actin-free XCAP1. Actin-free XCAP1 bound to both side and ends of actin filaments with moderate preference to ends. However, the XCAP1-actin complex bound preferentially to the side of actin filaments. These results indicate that binding of actin monomers to XCAP1 affects its binding modes to actin filaments, suggesting that this might be a novel mechanism to regulate the effects of CAP on actin filaments.

## Introduction

Both polymerization and depolymerization of actin are regulated in cells, and proper regulation of these processes are critical for cytoskeletal functions in many cell biological events (1-4). Among many actin-regulatory proteins, cyclase-associated protein (CAP) is a unique multi-functional regulator of actin filament dynamics (5, 6). CAP was originally discovered as a protein that binds to adenylate cyclase and is involved in cyclic AMP signaling in budding yeast (7, 8), and this function has been demonstrated more recently in mammalian cells (9). Moreover, CAP binds to both actin monomers and filaments to inhibit spontaneous polymerization (10, 11), enhance nucleotide exchange (12-17), promote filament severing (18, 19), and depolymerize filaments from pointed (20, 21) and barbed ends (22, 23). Physiological significance of the actin regulation by CAP has been supported by genetic studies showing that CAP knockout or knock down causes cytoskeletal abnormalities in many species including budding yeast (24), *Dictyostelium* (25), *Arabidopsis* (26), *Drosophila* (27, 28), *Caenorhabditis elegans* (15), and zebrafish (29). CAP is particularly important for the regulation of highly organized actin cytoskeleton in neurons (30-33) and striated muscles (15, 29, 34, 35). In humans, mutations in *CAP2*, encoding one of the two CAP isoforms, cause dilated cardiomyopathy and nemaline myopathy (36, 37). However, the effects of CAP on actin dynamics are complex, and how CAP exactly controls actin in cells remains largely unknown.

Three regions of CAP are known to bind to actin. The N-terminal helical-folded domain (HFD) binds to actin filaments on the side to enhance severing (18, 38) or at the pointed end to promote depolymerization (20, 21). However, whether the HFD binds to the side of actin filaments remains under debate, since one study reported negative results on the side binding (21). The central Wiskott Aldrich Syndrome Protein homology 2 (WH2) domain binds to actin monomers in competition with actin depolymerizing factor (ADF)/cofilin (17, 39) or to the barbed end of actin filament to induce depolymerization (22, 23). The C-terminal CAP and retinitis pigmentosa 2 (CARP) domain forms a dimer and binds to actin monomers to enhance nucleotide exchange (12, 40-44). Additionally, a putative coiled-coil at the very N-terminal end (45) and lateral association of HFD (46, 47) mediate CAP oligomerization, which enhances its activities to disassemble actin depolymerizing factor (ADF)/cofilin-bound actin filaments (18, 48, 49). Thus, actin binding to one of the three regions of CAP and/or its oligomerization states can affect the CAP activities, but interplays among these functional domains are not well understood.

Intriguingly, when CAP is purified from native sources, CAP is purified as an equimolar complex with G-actin (14, 19, 50-52). Stoichiometry of CAP and actin in the native or reconstituted complexes is reported as 6:6 (14, 38, 48) or 4:4 (51), and whether the difference in the stoichiometry is due to different species, isoforms, or experimental conditions are unknown. Importantly, the CAP-actin complex has activities to enhance turnover of ADF/cofilin-bound actin filaments (14), disassemble *Listeria* comet tails (19), and inhibit inverted formin 2 (INF2) (52), suggesting that the CAP-actin complex is a physiologically functional form of an actin regulator. However, to date, the functional significance of G-actin in the CAP-actin complex is not known. In this study, we used high-speed atomic force microscopy (HS-AFM) to characterize how actin-free *Xenopus* cyclase-associated protein 1 (XCAP1) or the XCAP1-actin complex interacts with actin filaments and found that actin monomers in the complex affect dwell time and positional preference of XCAP1 on actin filaments. Our results suggest that binding to actin monomers is a novel regulatory mechanism of CAP towards actin filaments.

## Results

### XCAP1 binds to actin filaments in a transient manner

In our previous study, we reported purification of a native complex of XCAP1 and G-actin at a 4:4 molar ratio (Fig. 1A and B, lane 2) (51). We found that G-actin could be removed by treating the complex with a high concentration of ATP, which is a much milder condition than the previously reported methods to dissociate G-actin from CAP, such as treatments with 3-4 M urea (12, 50). After the XCAP1-actin complex was adsorbed to a hydroxyapatite column, it was washed extensively with F-buffer containing 1 mM ATP to remove G-actin. Then, actin-free XCAP1 was eluted by a phosphate gradient (Fig. 1A and B lane 3). We used these preparations of XCAP1-actin complex and actin-free XCAP1 to characterize their interactions with F-actin using HS-AFM.

**Fig. 1.**
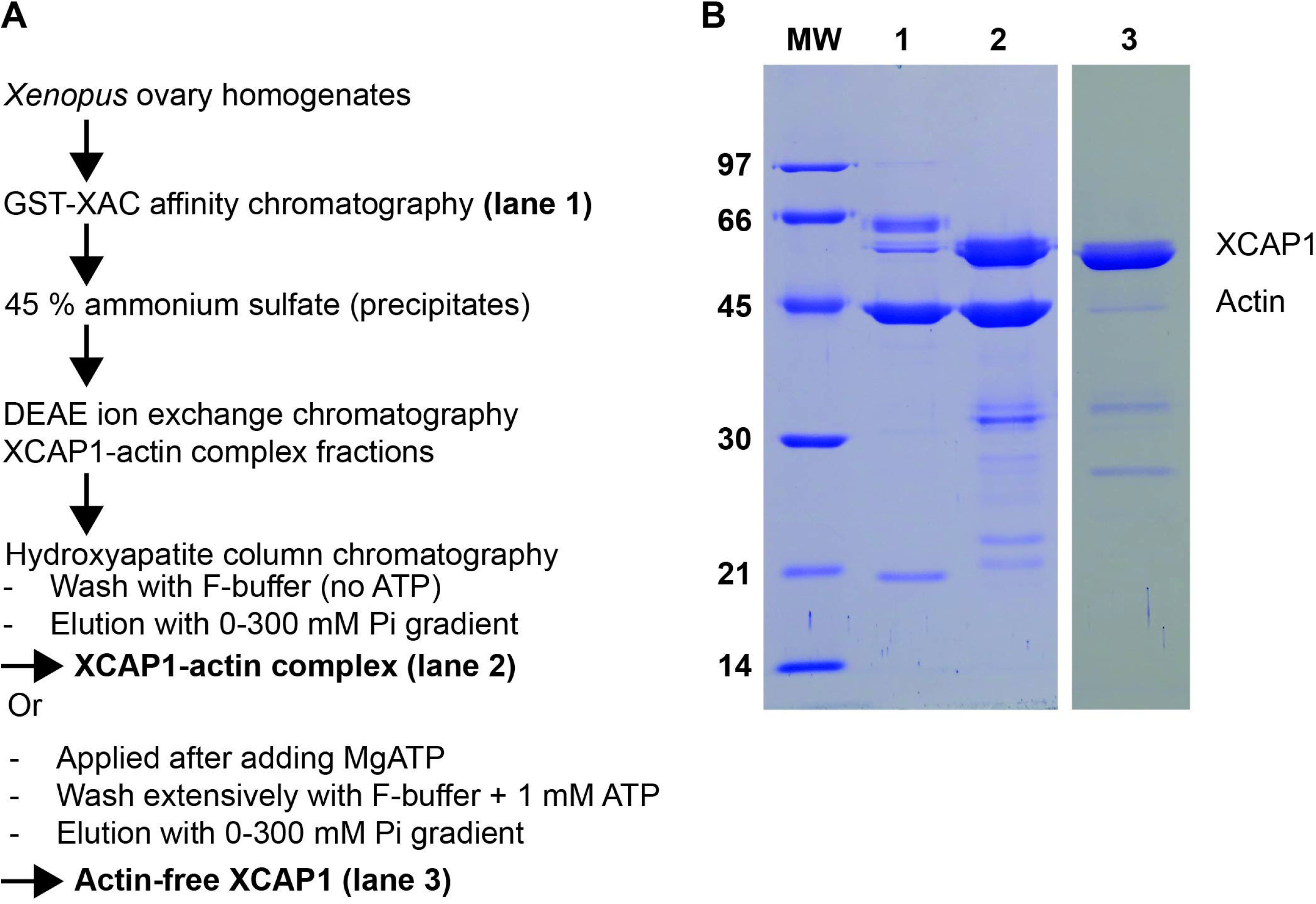
Purification of the XCAP1-actin complex and actin-free XCAP1 from *Xenopus laevis* oocytes. (A) Summary of the procedures to purify the XCAP1-actin complex and actin-free XCAP1. Details are described in Experimental Procedures. (B) Eluates from the GST-XAC affinity chromatography (lane 1), purified XCAP1-actin complex (lane 2), and purified actin-free XCAP1 (lane 3) were analyzed by SDS-PAGE and staining with Coomassie Brilliant Blue. Sizes of molecular weight markers (MW in kg/mol) are indicated on the left. Positions of XCAP1 and actin are indicated on the right.

To observe interactions of XCAP1 with F-actin, F-actin was attached to a lipid bilayer containing 89 % neutral lipid, DPPC, 10 % positively charged lipid, DPTAP, and 1 % biotin-cap DPPE (for facilitating spreading of the lipid bilayer) (53). The soft environments of the lipid bilayer preserve flexibility of the immobilized filaments, thereby minimizing artificial effects of immobilization (54-57). When actin-free XCAP1 was added to F-actin, bright particles, corresponding to XCAP1, bound to the actin filaments at the side and ends but stayed only for a short period (Fig. 2A arrowheads, Supplementary Movie 1) with an average dwell time of 0.2 sec (Fig. 2D, E). Severing or depolymerization events were very rare under our experimental conditions for both actin-free XCAP1 and the XCAP1-actin complex, therefore these were not analyzed in this study. Both volume analysis (Fig 2B) and height analysis of the XCAP1-bound regions (Fig. 2C) indicated that XCAP1 was in a single population with an average size of 320 nm^3^ and a height of 12.3 nm at a bound region (as compared to 8 - 9 nm at an unbound region), suggesting that XCAP1 was consistently at a tetrameric state (51) under our experimental conditions.

**Fig. 2.**
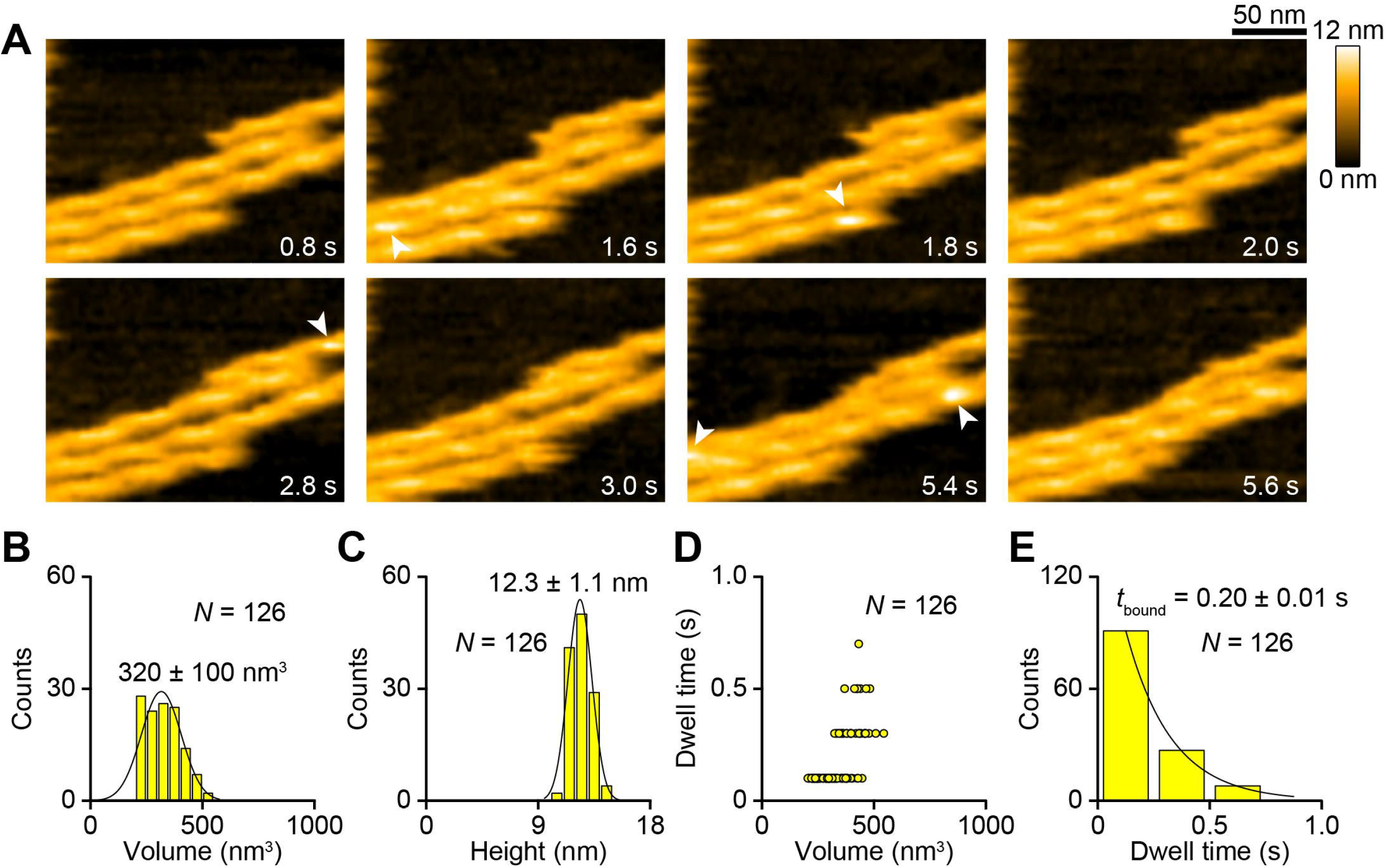
Actin-free XCAP1 binds to actin filaments in a transient manner. (A) Time-lapse HS-AFM images of actin-free XCAP1 interacting with F-actin on a lipid bilayer (see Supplementary Movie 1). Original scanning area was 300 × 150 nm^2^ with 200 × 50 pixels and cropped size was 200 × 150 nm^2^. Imaging rate was 0.2 s/frame (5 fps). Bar, 50 nm. Arrowheads indicate XCAP1 bound to F-actin. (B, C) Volume (B) and height (C) distributions of XCAP1 bound to F-actin. (D) Relationships between the volume and dwell time of XCAP1 on F-actin. (E) Dwell time analysis of XCAP1 on F-actin.

The XCAP1-actin complex also bound to the side of F-actin in a transient manner (Fig. 3A arrowheads, Supplementary Movie 2). However, unlike actin-free XCAP1, volumes of the actin-bound particles could be resolved into three sizes of peaks at 310, 510, and 720 nm^3^ (Fig. 3B), while the heights of the particle-bound regions were similar with an average of 12.3 nm (Fig. 3C). To understand the nature of the particles of different sizes, the molecular states of the XCAP1-actin complex were re-evaluated in the absence of F-actin on a mica surface rather than on a lipid bilayer. An advantage of using a mica surface is that a protein is strongly attached to the substrate allowing more precise estimation of the volumes of the protein. A disadvantage of using mica is that the strong adsorption of a protein complex to the substrate may promote partial disassembly of the complex (51). In the absence of F-actin, the XCAP1-actin complex could be resolved into at least six different sizes in the range from 1840 to 4000 nm^3^ (Supplementary Fig. 1). Note that these values are much larger than those of F-actin-bound XCAP1-actin complex (Fig. 3B), because flexible parts of the proteins on the lipid bilayer could not be resolved by HS-AFM (55, 58). Manual sorting of the images on the mica indicated that the XCAP1-actin complex was present in six distinct states with differences in brightness of the two lateral arm domains (Supplementary Fig. 1). Since each arm domain (the dimeric CARP domain) reversibly binds to two actin monomers (51), we predict that variable states of bound actin monomers in the complex caused the difference in the volumes of the complex (Fig. 3B).

**Fig. 3.**
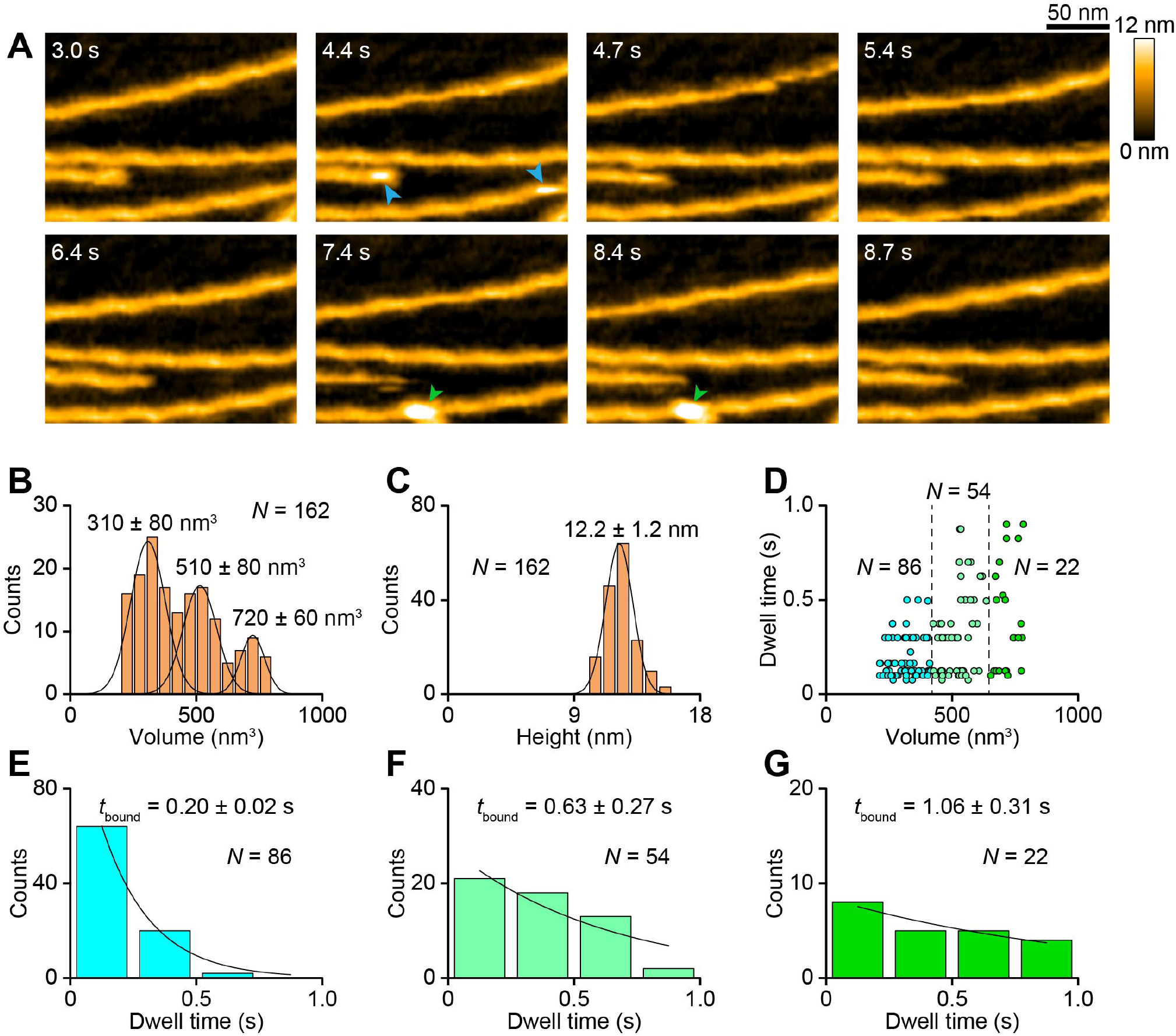
The XCAP1-actin complex binds to actin filaments in a transient manner. (A) Time-lapse HS-AFM images of the XCAP1-actin complex interacting with F-actin on a lipid bilayer (see Supplementary Movie 2). Original scanning area was 300 × 150 nm^2^ with 200 × 50 pixels and cropped size was 200 × 150 nm^2^. Imaging rate was 0.33 s/frame (3 fps). Bar, 50 nm. Arrowheads indicate the XCAP1-actin complex (blue: small; green: large) bound to F-actin. (B, C) Volume (B) and height (C) distributions of the XCAP1-actin complex bound to F-actin. The volume distribution indicates three major populations (small, medium, and large) of the XCAP1-actin complex on F-actin. (D) Relationships between the volume and dwell time of the XCAP1-actin complex on F-actin. Dashed lines at 417 and 680 nm^3^ indicate borders separating small, medium, and large particles. (E-G) Dwell time analyses of small (E), medium (F), and large (G) XCAP1-actin complex on F-actin.

Since the volume of the smallest peak matched with that of actin-free XCAP1 on F-actin (Fig. 2B), it represents XCAP1 with no bound G-actin. Because of the lower resolution of the volumes on F-actin, we assume that the medium and largest peaks corresponded to XCAP1 in complex with 1-2 and 3-4 actin monomers, respectively. Although the particles of all sizes bound to F-actin in a transient manner, there was a positive correlation between the dwell time and the particle volume (Fig. 3D). Dwell times of small (<417 nm^3^), medium (417-680 nm^3^), and large (>680 nm^3^) particles were separately determined as 0.20, 0.63, and 1.06 sec (Fig. 3E-G), respectively. The average dwell time of the small particles matched with that of actin-free XCAP1 (Fig. 2E), indicating again that the small particles represent XCAP1 with no bound G-actin. Therefore, the XCAP1-actin complex containing 3 or 4 actin monomers bound to F-actin 5-fold longer than XCAP1 alone, and the XCAP1-actin complex containing 1 or 2 actin monomers did 3-fold longer than XCAP1 alone. Thus, binding of G-actin to XCAP1 causes significant effects to prolong its F-actin binding.

### G-actin influences the preference of XCAP1 for side or end binding on F-actin

Next, we analyzed positional preference of actin-free XCAP1 or the XCAP1-actin complex for F-actin binding. For this purpose, we selected actin filaments only when at least one end was included throughout time-lapse imaging and analyzed end- or side-binding of XCAP1 or the XCAP1-actin complex. Note that we could not define the polarity of the filaments in our experiments. Therefore, the end binding was either at the pointed (20, 21) or barbed end (22, 23). Both actin-free XCAP1 (Fig. 4A, B) and the XCAP1-actin complex (Fig. 5A-C) bound to ends and sides. In typical images, binding sites on the filament sides are present at a much larger number than those at filament ends. Therefore, simple counting of binding positions can result in a large number of side-binding events (Fig. 4C and 5D) and may not be good parameters to determine a positional preference for XCAP1 binding. To overcome this problem, we calculated occupancy ratios for end or side as described in Experimental Procedures. This analysis showed that actin-free XCAP1 occupied end more frequently than side (Fig. 4D). Dwell time of actin-free XCAP1 at the end (0.24 ± 0.064 sec; Fig. 4E) was similar to that at the side (0.20 ± 0.012 sec; Fig. 4F). Similarly, the small particles in the XCAP1-actin complex (XCAP1 with no bound G-actin) occupied end more frequently than side (Fig. 5E). However, the medium particles in the XCAP1-actin complex occupied side more frequently than end, and the large particles bound only to the side (Fig. 5E). These results indicate that, in the absence of G-actin, XCAP1 binds preferentially to filament ends, whereas binding of G-actin to XCAP1 alters its preference to side-binding.

**Fig. 4.**
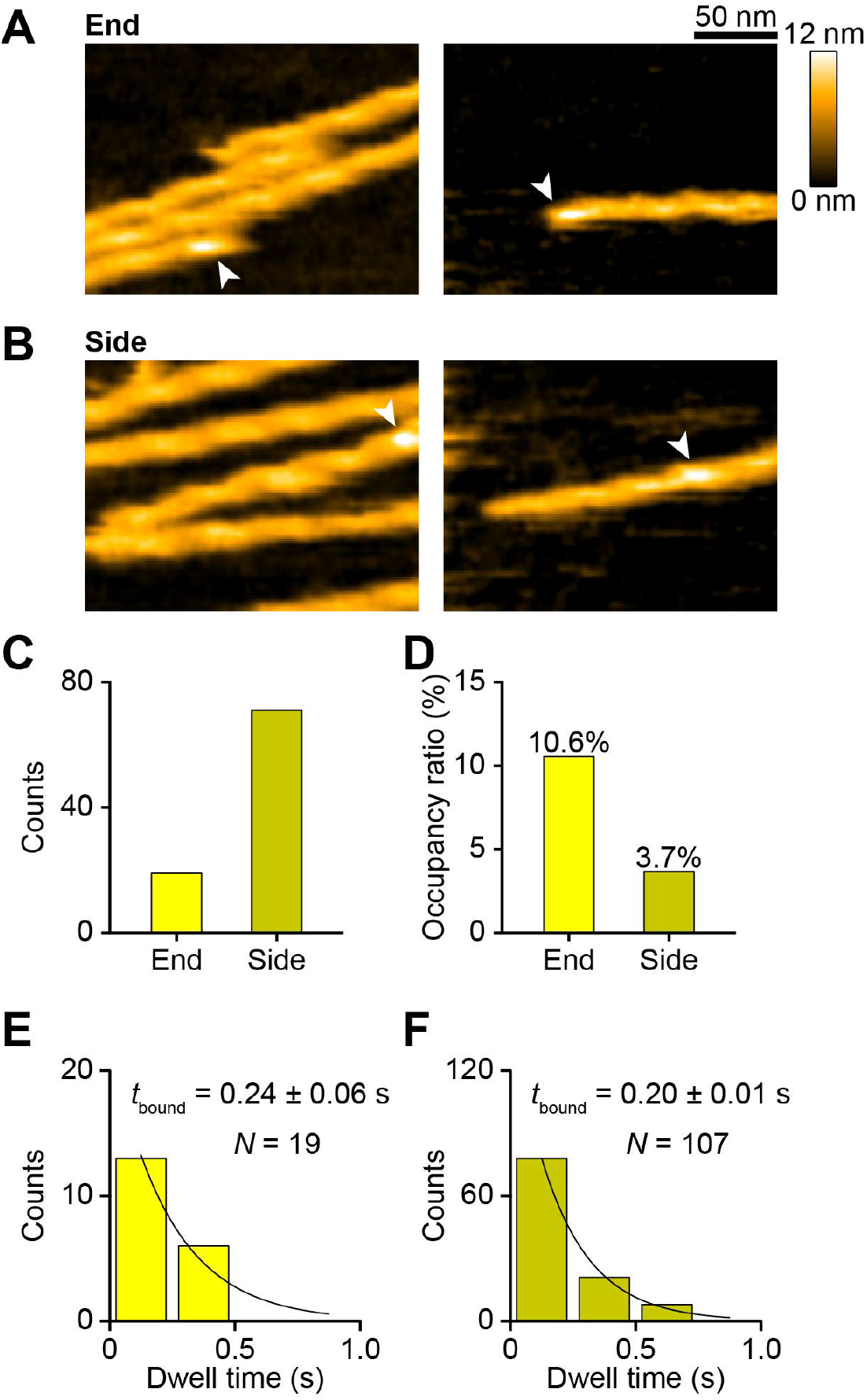
Actin-free XCAP1 binds to F-actin with a moderate preference for ends. (A, B) Representative HS-AFM images of actin-free XCAP1 bound to end (A) or side (B) of F-actin as indicated by arrowheads. Original scanning area was 300 × 150 nm^2^ with 200 × 50 pixels (left) or 300 × 180 nm^2^ with 200 × 60 pixels (right). Cropped size was 200 × 150 nm^2^ (left) or 200 × 150 nm^2^ (right). Imaging rate was 0.2 s/frame (5 fps). (C, D) Counts (C) and occupancy ratios (D) of actin-free XCAP1 bound to end or side. (E, F) Dwell time analyses of actin-free XCAP1 bound to end (E) or side (F) of F-actin.

**Fig. 5.**
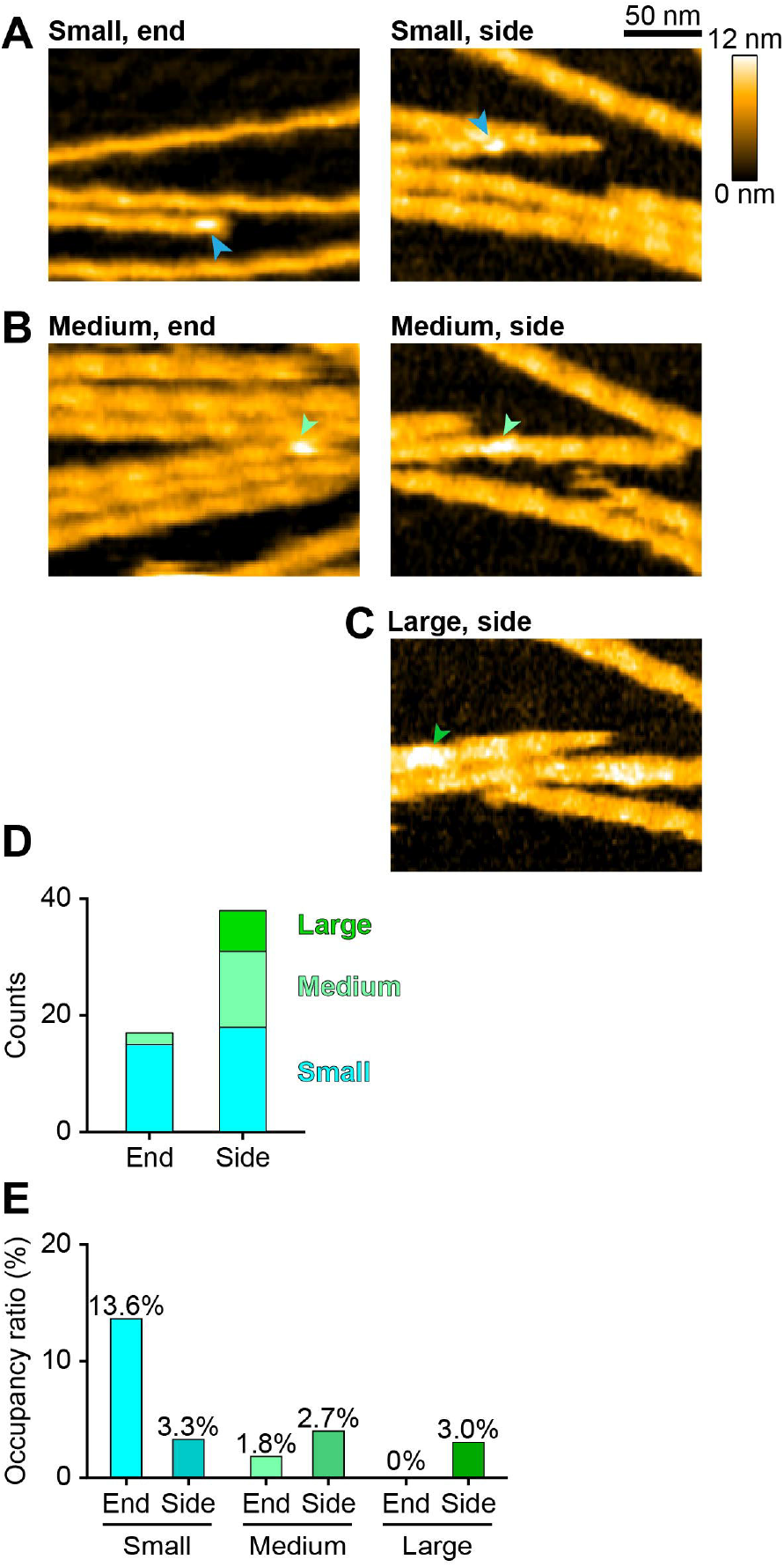
G-actin alters positional preference of XCAP1 on F-actin. (A-C) Representative HS-AFM images of the small (A), medium (B), or large (C) XCAP1-actin complex bound to end (left) or side (right) of F-actin as indicated by arrowheads. Original scanning area was 300 × 150 nm^2^ with 200 × 50 pixels (A, left), 400 × 200 nm^2^ with 200 × 50 pixels (B, left), or 300 × 180 nm^2^ with 200 × 60 pixels (A-C, right). Cropped size was 200 × 150 nm^2^ for all images. Imaging rate was 0.33 s/frame (3 fps) (A, left), 0.4 s/frame (2.5 fps) (B, left), or 0.25 s/frame (4 fps) (A-C, right). (D, E) Counts (D) and occupancy ratios (E) of the XCAP1-actin complex (small: blue; medium: light green; large: green) bound to end or side of F-actin.

## Discussion

In this study, we used HS-AFM and demonstrated that XCAP1 bound to sides and ends of F-actin in a transient manner. Both actin-free XCAP1 and XCAP1 in complex with G-actin bound to F-actin, but the bound G-actin influenced the dwell time and positional preference of XCAP1 on F-actin (Fig. 6). The XCAP1-actin complex bound to F-actin 3-to 5-fold longer than actin-free XCAP1. Actin-free XCAP1 bound preferentially to filament ends, whereas the XCAP1-actin complex did to filament side. Previous studies have shown that CAP binds to the pointed ends (20, 21), barbed ends (22, 23), and side of F-actin (18, 19, 38). However, the study by Kotila et al. (21) reported that the N-terminal HFD of mouse CAP1 (N-CAP) did not bind to the side of F-actin and could not be docked structurally to the side of F-actin. One possible reason for the discrepancy is the transient nature of interactions that may not be captured reliably by fluorescent imaging. Our HS-AFM system was suitable to characterize transient CAP-F-actin interactions. In addition, CAPs from different species and/or different sources may exhibit different F-actin binding properties. We used XCAP1 and the XCAP1-actin complex from a non-recombinant native source with no additional labeling, which should have minimized artificial effects.

**Fig. 6.**
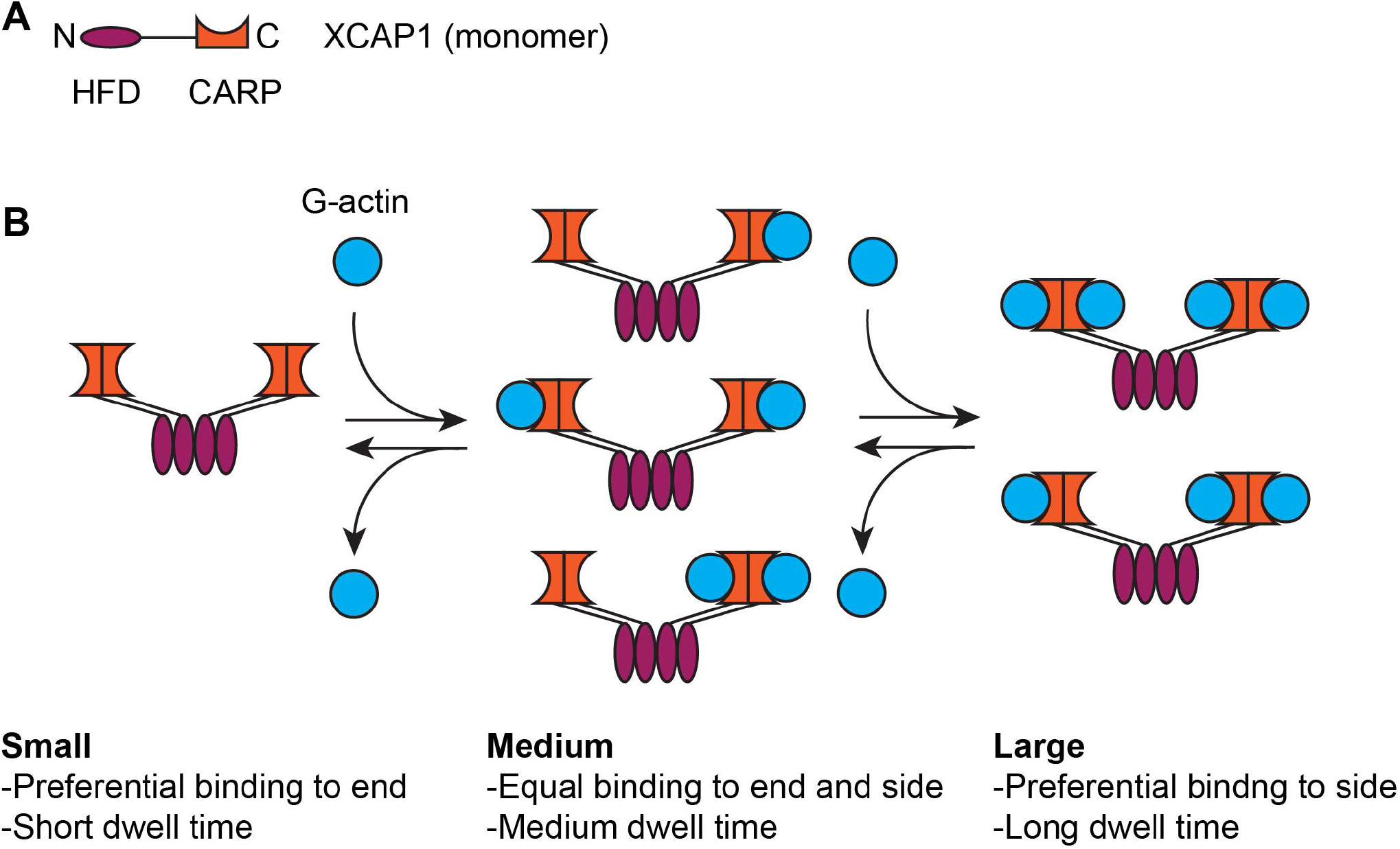
Summary of the effects of G-actin bound to XCAP1 on F-actin binding. (A) A simplified model of an XCAP1 monomer (N- and C-termini are indicated by N and C) only shows the HFD and CARP domains. (B) G-actin reversibly interacts with CARP of the XCAP1 tetramer (51) and influences positional preference and dwell time of XCAP1 on F-actin.

As mentioned above, a role of the N-terminal HFD of CAP in binding to the side of F-actin remains controversial. Alternatively, we consider the possibility that the WH2 domain of CAP may participate in side binding. The WH2 domain of CAP was previously shown to bind to G-actin and be important for promoting exchange of G-actin-bound nucleotides (17, 39). It is also a critical domain to promote barbed-end depolymerization (22, 23). A peptide containing the WH2 domain of mouse CAP2 promotes severing of actin filaments (35), and the WH2 domain of yeast Srv2/CAP is required for bundling actin filaments (59), suggesting that the WH2 of CAP directly binds to the side of actin filaments. Similarly, WH2 domains of other actin-regulatory proteins including Cordon-Bleu (60, 61), Spire (62, 63), and INF2 (64, 65) also interact with the side of actin filaments. Since G-actin affects the dwell time and positional preference of XCAP1 on F-actin, we hypothesize that G-actin bound to the C-terminal CARP domain may alter the accessibility of WH2 to the side or ends of F-actin, thereby influencing the interaction between XCAP1 and F-actin. However, further biochemical and structural studies are required to understand this mechanism.

Complex formation between CAP and G-actin is conserved across eukaryotes, and previous studies on the CAP-G-actin interaction primarily focused on the effects on monomer sequestration and nucleotide exchange (5, 6). However, our results indicate a novel aspect of the effects of G-actin on the interaction between CAP and F-actin (Fig. 6). G-actin-free XCAP1 bound preferentially to the filament ends, but the XCAP1-actin complex bound to the side, suggesting that G-actin-free CAP is a suitable form to depolymerize actin filaments from pointed or barbed ends. Therefore, G-actin could be an inhibitor of the end depolymerase activity of CAP. Unfortunately, we have not been able to reconstitute end depolymerization by XCAP1 in the presence or absence of cofilin in HS-AFM and could not test this hypothesis (our unpublished observations). If this is correct, G-actin binding to the CARP domain of CAP could be a regulatory mechanism for filament depolymerization. The CARP domain of CAP binds preferentially to ADP-actin over ATP-actin (17, 40). We also showed that excess ATP dissociates G-actin from CAP (Fig. 1). Thus, when ATP is constantly generated in cells, ADP-actin is converted to ATP-actin and dissociated from the CARP domain. However, when ATP is depleted, ADP-actin will make a tight complex with CAP and may inhibit its end depolymerization activity.

The transient nature of F-actin binding by XCAP1 may explain how CAP-induced F-actin bundles are remodeled in a dynamic manner. Yeast Srv2/CAP alone or together with actin-binding protein 1 bundles actin filaments and allows filament sliding to maximize filament overlaps, which results in remodeling of loose actin filaments into compact thick actin bundles (59). In the case of Srv2/CAP alone, the WH2 domain is required for the bundling activity (59). If Srv2/CAP tightly binds to the side of F-actin, the filaments will not be allowed to slide and remodel. Therefore, a short dwell time of CAP on F-actin should allow cycles of association and dissociation until the actin bundles reach an energetically stable compact state.

In conclusion, our study revealed a new regulatory mechanism of the CAP-F-actin interaction by G-actin, indicating that the functional relationship between CAP and G-/F-actin is more complex than previously thought. It will also be important to clarify how ADF/cofilin participates in their interactions. CAP and ADF/cofilin synergistically promote filament severing (18, 19) and pointed-end depolymerization (20, 21). Additionally, CAP and ADF/cofilin compete for G-actin binding (12, 16, 40), but how this competition on G-actin influences filament severing and depolymerization is unknown. Furthermore, phosphoregulation of CAP (66, 67) and ADF/cofilin (68, 69) may affect the balance of interactions among CAP, ADF/cofilin, and actin during rearrangement of the actin cytoskeleton in cells.

## Experimental Procedures

### Actin preparation

Rabbit skeletal muscle actin was purified as described previously (70). Purified G-actin (∼50 μM) was rapidly frozen in liquid nitrogen in aliquots of 10 μl and stored at −80 °C. Prior to HS-AFM experiments, an aliquot was rapidly thawed by hands and stored on ice. Actin filaments (F-actin) of 10 μM based on the G-actin concentration were formed by adding final concentrations of 100 mM KCl, 2 mM MgCl_2_, 1 mM EGTA and 1 mM ATP at room temperature for one hour. Then, F-actin was stored on ice until use and used within three days.

### Purification of XCAP1-actin complex and actin-free XCAP1 from Xenopus laevis oocytes

XCAP1-actin complex was purified from *Xenopus laevis* oocytes as described previously (51). From this procedure, the DE52 ion-exchange chromatography fractions containing pure XCAP1-actin complex were further processed to obtain actin-free XCAP1 (Fig. 1). After adding 2 mM MgCl_2_ and 1 mM ATP to the pooled fractions, these were applied to a hydroxyapatite column preequilibrated with an F-buffer (0.1 M KCl, 2 mM MgCl_2_, 0.1 mM DTT, 0.01% NaN_3_, and 20 mM HEPES-KOH, pH 7.2) containing 1 mM ATP, and thoroughly washed with the same buffer to remove actin. Then, XCAP1 was eluted using a linear gradient of 0 to 300 mM potassium phosphate buffer at pH 7.2 in F-buffer. Purified XCAP1 was concentrated by ultrafiltration with Ultracel-30K (Millipore) and dialyzed against an F-buffer.

### High-speed atomic force microscopy

High-speed atomic force microscopy (HS-AFM) imaging was performed in aqueous buffer at room temperature using a laboratory-built setup, as described previously (53). A glass sample stage (2 mm diameter, 2 mm height) was fixed to the Z-scanner by nail polish, with a thin mica disc (1.5 mm diameter, ∼0.05 mm thickness) affixed to the top via epoxy. Samples were deposited on either bare mica or mica-supported lipid bilayers. Imaging was performed in tapping mode using small cantilevers (BLAC10DS-A2, Olympus, Japan) with a resonant frequency of ∼0.5 MHz in water, a quality factor of ∼1.5 in water, and a spring constant of ∼0.1 N/m. To increase the spatial resolution, an additional probe tip of ∼500 nm long was grown on the original tip end of a cantilever through electron beam deposition using a field-emission scanning electron microscope (Verious 5UC, Thermo Fisher Scientific, USA). Ferrocene was used as a gas source. The additional probe tip was further sharpened using a radio-frequency plasma etcher (Tergeo, PIE Scientific LLC, USA) under an argon gas atmosphere (Direct mode, 10 sccm, and 20 W for 1.5 min). For HS-AFM imaging, the cantilever’s free-oscillation peak-to-peak amplitude (*A*_0_) was set at approximately 1.5-2.0 nm, with a feedback amplitude set-point of ∼0.9×*A*_0_. To observe the interactions between actin filaments and XCAP1, we used the only tracing imaging (OTI) mode (71) for reducing the tip-sample interaction force while increasing the scanning speed. Imaging parameters, imaging rate, scan area, and pixel resolution are detailed in each figure legend.

### Mica-supported lipid bilayers

Small unilamellar vesicles (SUVs) and mica-supported lipid bilayers were prepared, as described previously (53, 72). Briefly, lipids of 1,2-dipalmitoyl-*sn*-glycero-3-phosphocholine (DPPC), 1,2-dipalmitoyl-3-trimethylammonium-propane (DPTAP), and 1,2-dipalmitoyl-*sn*-glycero-3-phosphoethanolamine-*N*-(cap biotinyl) (biotin-Cap-DPPE) were purchased from Avanti Research (Birmingham, Alabama). A lipid composition of DPPC:DPTAP:biotin-cap-DPPE at 89:10:1 (w/w/w) was used. The liposome solution dissolved in Milli-Q water at 2□mg/ml was prepared and stored as 10□μl aliquots at −20□°C before use. After thawing an aliquot, the liposome solution was diluted to 0.5 mg/ml with 10 mM MgCl_2_ and sonicated for 3 min in a bath sonicator (AUC-06L, AS ONE, Japan) to produce SUVs. The SUVs solution of 2 µl was applied to freshly cleaved mica and incubated for at least 3 hours at room temperature (24–26 °C) in a humidified, sealed chamber. This promoted lipid bilayer formation and prevented drying. Up to ten stages with mica-supported lipid bilayer were prepared in parallel under identical conditions and stored before use.

### HS-AFM imaging

To observe the molecular features of the XCAP1-actin complex, a drop (2 μl) of diluted protein sample (ca. 1 nM) with buffer A (100 mM KCl, 2 mM MgCl_2_, 20 mM HEPES–KOH, pH 7.2) was deposited on a mica surface for 5 minutes. Then, the surface was rinsed with 20 μl of buffer A and imaged in buffer A of 60 μl containing 100 nM ADP-G-actin.

To observe the interactions between actin filaments (F-actin) and XCAP1-actin complex or actin-free XCAP1, the mica-supported lipid bilayer was used. Before F-actin deposition, the surface of the sample stage with mica-supported lipid bilayer was rinsed with Milli-Q water (20 µl ×5) to remove excess SUVs. Water on the sample stage was replaced with 2 µl of the observation buffer (30 mM KCl, 10 mM imidazole-HCl, pH 7.0, 1 mM MgCl_2_, 0.5 mM EGTA, 10 mM DTT, 10 % glycerol, 0.2 mM ATP). A drop of 2 µl F-actin diluted to 1 µM with the observation buffer was applied to the sample stage. After incubation for 10 minutes at RT, unbound filaments were washed using the observation buffer. The sample stage was then immersed in the observation buffer of 60 μl for imaging. To observe binding dynamics of actin-free XCAP1 or the XCAP1-actin complex, 6 µl of the observation buffer containing either actin-free XCAP1 or the XCAP1-actin complex was injected into the liquid chamber during AFM imaging at final concentration of 20 – 100 nM.

### HS-AFM data analysis

AFM data were processed and quantified using a laboratory-made UMEX Viewer and UMEX Viewer for Height Analysis, as described in previous studies (57). Each raw image was first filtered with a low-pass filter to remove spike noise and with a flattening filter to make the xy-plane flat.

The molecular height was measured by calculating the difference between the maximum height of the molecule and the average height of the substrate. For volume measurements, the area occupied by a target molecule was first recognized by comparing the average height of the substrate. Then, the volume was calculated by accumulating the height containing each pixel within that area. Binding lifetimes of actin-free XCAP1 or the XCAP1-actin complex on F-actin were obtained by measuring the dwell time of individual molecules. AFM images and movies were processed in UMEX Viewer and After Effects (Adobe, USA), and all statistical evaluations were carried out in OriginPro 2024 (OriginLab Corporation, USA).

### Calculation of the occupancy ratios for binding sites

From the acquired HS-AFM images, images of actin filaments showing at least one end were selected and processed for analysis. Total length of the observed F-actin (*L*_a_) was measured, and the number of F-actin with binding event (*N*_a_) was counted. The number of protomers that can make the side binding (assuming that one actin protomer provides one binding site) for actin-free XCAP1 or XCAP1-actin complex in the i-th F-actin *N*_s-i_ was calculated as (*N*_s-i_ = 2*L*_a_/*D*_aa_ – *N*_e_), where *D*_aa_ (= 5.5 nm) and *N*_e_ (= 2) are the distance between adjacent protomers and the number of protomers at the end of F-actin, respectively. However, due to steric hindrance by the substrate surface, a half of protomers were considered inaccessible to actin-free XCAP1 or XCAP1-actin complex. Thus, only the protomers of *N*_s-i-eff_ (= *L*_a_/*D*_aa_ – *N*_e_) were effectively available for binding to actin-free XCAP1 or XCAP1-actin complex. The occupancy ratios for each binding site (end or side) were determined by:

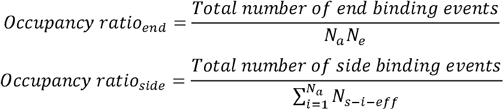

## Abbreviations

ADF: (actin depolymerizing factor)
CAP: (cyclase-associated protein)
CARP: (CAP and retinitis pigmentosa 2)
HFD: (helical-folded domain)
HS-AFM: (high-speed atomic force microscopy)
WH2: (Wiskott Aldrich Syndrome Protein homology 2)
XCAP1: (*Xenopus* cyclase-associated protein 1)

## Acknowledgements

We thank Prof. Toshio Ando, Dr. Kenichi Umeda, Ms. Kayo Nakatani and Ms. Risa Omura for technical support of the HS-AFM experiments. This work was supported by grants from the National Institutes of Health (R01-GM144563) to SO and the World Premier International Research Center Initiative (WPI), MEXT, Japan, KAKENHI (24H00402 to NK), JSPS, Japan.

## Data availability

All data are contained in the article.

## Supporting Information

**Supplementary Figure 1. Classification of different molecular states of the XCAP-actin complex**. Representative HS-AFM images (A), volume distributions (B), and models (C) of six different states of the XCAP1-actin complex on mica surfaces. Bar, 20 nm. Original scanning area was 150 × 150 nm^2^ with 80 × 80 pixels, and cropped size was 60 × 60 nm^2^. Imaging rate was 0.2 s/frame (5 fps). The images show that they are different only in the lateral arm domains representing the CARP domain of XCAP1 that reversibly interacts with G-actin (ref. 51).

**Supplementary Movie 1. Time-lapse HS-AFM movie of actin-free XCAP1 interacting with F-actin on a lipid bilayer**. Scanning area was 300 × 150 nm^2^ with 200 × 50 pixels. Imaging rate was 0.2 s/frame (5 fps). Arrowheads indicate XCAP1 bound to F-actin.

**Supplementary Movie 2. Time-lapse HS-AFM movie of the XCAP1-actin complex interacting with F-actin on a lipid bilayer**. Scanning area was 300 × 150 nm^2^ with 200 × 50 pixels. Imaging rate was 0.33 s/frame (3 fps). Arrowheads indicate the XCAP1-actin complex (blue: small; green: large) bound to F-actin.

