## Supplementary Fig. 1 for "Actin monomers influence the interaction between *Xenopus* cyclase-associated protein 1 and actin filaments": Supplementary Fig1.pdf

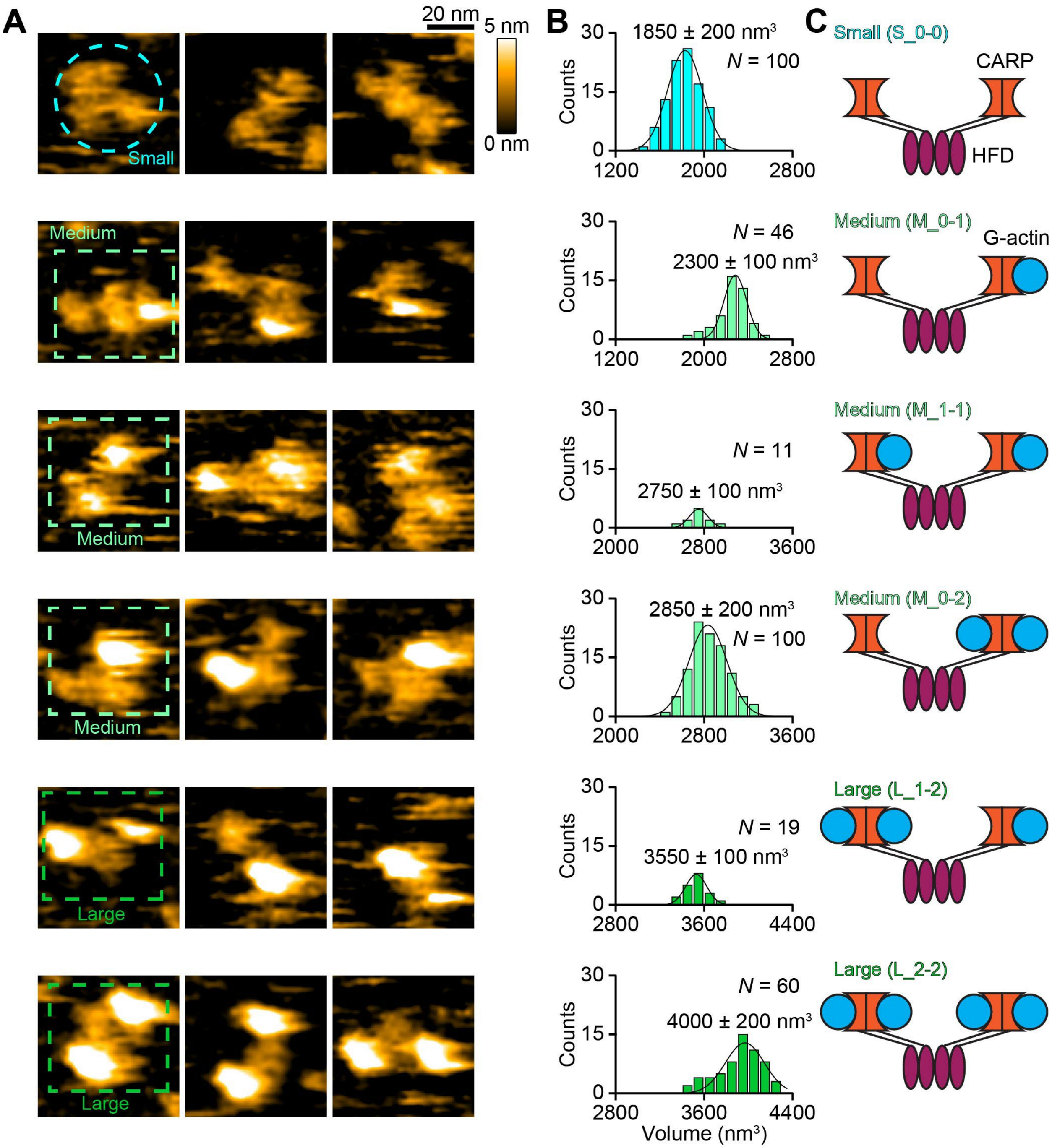


**Supplementary Figure 1. Classification of different molecular states of the XCAP-actin complex.** Representative HS-AFM images (A), volume distributions (B), and models (C) of six different states of the XCAP1-actin complex on mica surfaces. Bar, 20 nm. Original scanning area was 150 × 150 $nm^2$ with 80 × 80 pixels, and cropped size was 60 × 60 $nm^2$. Imaging rate was 0.2 s/frame (5 fps). The images show that they are different only in the lateral arm domains representing the CARP domain of XCAP1 that reversibly interacts with G-actin (ref. 51).
